# IL-34 empowers regulatory T cells with novel non-canonical function to safeguard brain barrier integrity during neuro-inflammation

**DOI:** 10.1101/2024.09.09.611994

**Authors:** Lien Van Hoecke, Janne Verreycken, Lore Van Acker, Laura Amelinck, Junhua Xie, Jonas Castelein, Elien Van Wonterghem, Griet Van Imschoot, Marlies Burgelman, Sarah Vanherle, Ilse Dewachter, Paulien Baeten, Bieke Broux, Roosmarijn E. Vandenbroucke

## Abstract

In efforts to find reparative strategies for brain damage, brain-associated regulatory T cells (Tregs) have gained increasing attention in recent years. Beyond their textbook immunoregulatory function, Tregs have emerged as key players in the response to brain trauma and the restoration of damaged brain tissue. Here, we are the first to describe a novel, non-canonical function of Tregs in maintaining the sealing capacity of both the blood-brain barrier (BBB) and the blood-cerebrospinal fluid (CSF) barrier. Moreover, we identified the cytokine IL-34 as a critical determinant in this newly unveiled Treg function. Mechanistically, IL-34 exerts its influence by modulating the expression and localization of the tight junction protein ZO-1 in both BBB endothelial cells and choroid plexus epithelial cells, thereby reinforcing the strength of the brain barriers. Given the well-established notion of leaky brain barriers and the involvement of immunological components in neurological diseases such as Alzheimer’s disease (AD) and multiple sclerosis (MS), we further demonstrate diminished IL-34 expression in Tregs derived from patients with relapsing-remitting MS (RR-MS) and patients with AD and even mild cognitive impairment (MCI). Remarkably, our study reveals the potential of IL-34 treatment in reinstating the integrity of brain barriers within murine models mimicking these neurological disorders. These ground-breaking findings shed light on the intricate relationship between Tregs, IL-34, and the integrity of brain barriers. They offer novel avenues for therapeutic approaches to ameliorate brain barrier dysfunction in the context of neurological disorders.

## Introduction

In recent years, there has been a notable shift away from the concept of brain immune privilege^1–4^, as we now acknowledge the presence of immune cells within the central nervous system (CNS) and their active involvement in immune responses^5–9^. This shift in understanding has also transformed the way we perceive the pathology of neurological disorders, transitioning from a ’neuro-centric’ perspective to a more ’inflammation-centric’ viewpoint. The presence of immune cells in the CNS has dual implications. On one hand, it can be advantageous as these immune cells clear debris and facilitate injury repair. However, the flip side of this scenario emerges when inflammation becomes persistent and chronic, leading to neuronal dysfunction, injury, and ultimately, neurodegeneration. Remarkably, chronic neuro-inflammation seems to be a common element in most, if not all, neurological disorders. Consequently, the field of immunotherapy for neurological disorders is rapidly expanding, with increasing attention being given to regulatory T cells (Tregs)^10,11^.

Beyond their well-established role in immune system regulation, Tregs are now also recognized for their significant contributions to maintaining tissue balance and promoting repair in various models of neurological disorders^12–16^. Due to these unique attributes, therapies aimed at enhancing Treg activity hold tremendous promise as a potential ’holy grail’ for the treatment of inflammatory neurological disorders characterized by excessive neuro-inflammation and impaired brain repair.

Next to immune cells in the brain parenchyma, it is also evident that immune cells positioned at different brain borders, such as the meninges, choroid plexus (ChP), and perivascular spaces at the blood-brain barrier (BBB), play a crucial role in immune surveillance^17–20^. These compartments serve as sites where diverse immune cells sample CNS antigens, protect the brain from pathogenic insults, influence behavioural paradigms, and contribute to immune cell migration to the brain^8,9,21–24^. Increased inflammation at the brain barriers is known to result in loss of tight junctions (TJs) and adherens junctions (AJs), thereby opening the gateway to the CNS parenchyma. Importantly, the breakdown of these barriers contributes to various neurological disorders, such as multiple sclerosis (MS), stroke, brain trauma, and Alzheimer’s disease (AD)^25–27^. As a consequence, the loss of barrier integrity results in infiltration of immune cells, and disruption of transport and clearance of molecules, thereby contributing to disease progression^2^. Consequently, repairing brain barrier integrity in neurological disorders represents a promising therapeutic avenue.

Research on the dynamic relationship between brain barriers and the immune system has primarily focused on understanding how immune cells breach these protective barriers to enter the parenchyma. Yet, the exploration into how immune cells influence the sealing function of these barriers remains considerably understudied. In this study, we have identified a novel Treg-brain barrier interaction that holds tremendous potential for treating patients with neurodegenerative disorders. Through transfer and depletion experiments, we have pinpointed the crucial involvement of Tregs in fortifying both the BBB and the blood-CSF barrier. Mechanistically, we have identified the cytokine IL-34 as a key factor in this newfound function of Tregs. Specifically, IL-34 increases the expression and continuous fragment length of the TJ protein ZO-1 in BBB endothelial cells (BBB-EC) and ChP epithelial (CPE) cells, thereby enhancing the strength of the brain barriers. Interestingly, we have observed reduced IL-34 expression in Tregs from patients with relapsing-remitting multiple sclerosis (RR-MS) and AD. We even identified a decreased percentage of IL-34 expressing Tregs in persons with mild cognitive impairment (MCI), which is an early sign of dementia, suggesting that this mechanism is involved in neurodegeneration from the start of pathology. In terms of the therapeutic implications of our findings, we present compelling evidence showcasing the efficacy of IL-34 treatment in reinstating the integrity of brain barriers in murine models that accurately mimic these neurological disorders.

In summary, our findings reveal that IL-34 orchestrates a novel non-canonical function of Tregs during neuro-inflammation by preserving the sealing function of both the BBB and the blood-CSF barrier. These discoveries shed light on the intricate interplay between Tregs, IL-34, and brain barriers, offering promising avenues for therapeutic strategies aimed at alleviating brain barrier impairments in neurological disorders.

## Results

### Enhanced brain barrier permeability and impaired tight junction ZO-1 expression in mice lacking a mature adaptive immune system

To investigate the impact of the adaptive immune system on blood-brain barrier (BBB) and blood-cerebrospinal fluid (CSF) barrier integrity, we conducted a comparative analysis between wild type mice and *Rag2*^-/-^ mice, i.e. mice lacking mature T and B lymphocytes. In mice lacking a functional adaptive immune system, compared to control mice, there was increased leakage of 4 kDa dextran in various regions of the brain parenchyma (including the hippocampus, cortex, and cerebellum) as well as in the CSF upon systemic administration, representative of increased BBB and blood-CSF barrier leakage, respectively (**Fig. 1A** and **1C**). This observation was confirmed via confocal imaging visualizing 4 kDa biotin dextran (BD) in the brain parenchyma and at the CSF side of the choroid plexus (ChP) of *Rag2*^-/-^ mice systemically injected with BD (**Fig. 1B** and **1D**).

**Figure 1.**
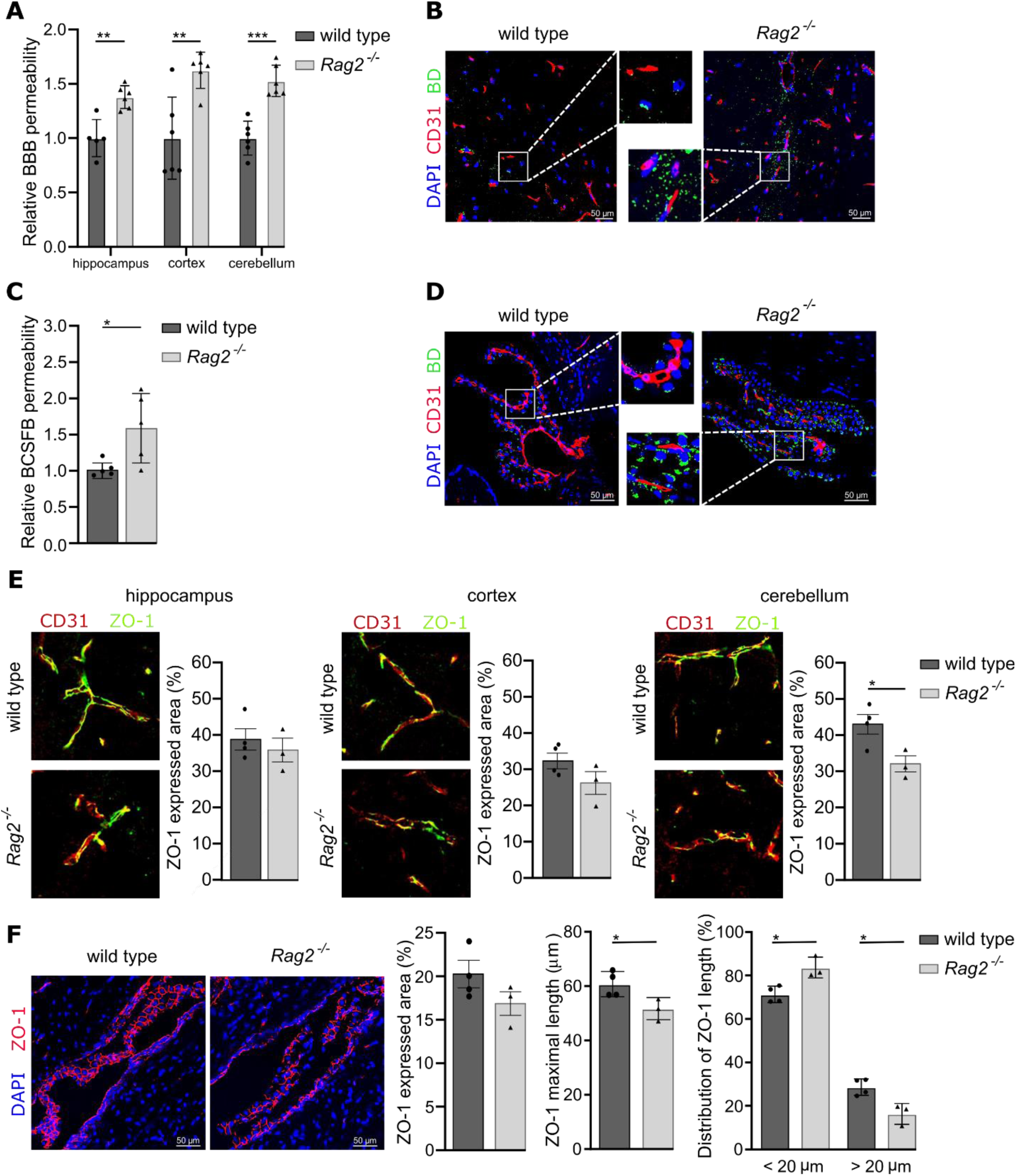
The barrier integrity of the blood-brain barrier (BBB) and blood-cerebrospinal fluid (CSF) barrier is compromised in the absence of a mature adaptive immune system. **(A, C)** Barrier permeability was determined 15 min after intravenous injection of 4 kDa FITC-dextran for the BBB (A) and the blood-CSF barrier (C). Graphs represent relative permeability normalized to wild type mice (n=5-6). **(B, D)** Representative images of immunohistochemical staining for biotin-dextran (BD; green), in combination with an endothelial marker (CD31; red) at the BBB (B) and blood-CSF barrier (D). **(E-F)** Representative images and quantification of staining for tight junction protein Zonula occludens-1 (ZO-1), in combination with CD31 staining at BBB (E) and at blood-CSF barrier (F). The graphs are shown as the mean ± SEM and the datapoints are biological replicates. Images are representative for 4 biological replicates. Statistical analysis was performed with Mann-Whitney test. * *p* < 0.05; ** *p* < 0.01; *** *p* < 0.001.

Confocal imaging further demonstrated a more diffuse distribution of TJ protein ZO-1 in *Rag2^-/-^* mice compared to their wild type counterparts at the BBB (**Fig. 1E**) and at the blood-CSF barrier (**Fig. 1F**). Notably, a significant reduction in ZO-1 expression was observed at the BBB in the cerebellum of *Rag2*^-/-^ mice compared to wild type mice (**Fig. 1E**), and a trend towards reduction at the blood-CSF barrier (**Fig. 1F**). Moreover, when analysing the maximal length of continuous ZO-1 subcellular localisation, a significant reduction in ZO-1 fragments longer than 20 µm could be observed in *Rag2^-/-^* compared to wild type mice at the blood-CSF barrier, indicating a difference in distribution of continuous ZO-1 signal.

Together, these findings reveal that the adaptive immune system plays a crucial role in the maintenance of brain barrier integrity.

### Novel non-canonical function of Tregs as key player in the formation of tight brain barriers

The effects of the overall adaptive immune system on the sealing capacity of the brain barriers in *Rag2^-/-^* mice can be attributed to T cells, B cells, or a combination of both lymphocyte populations. Transfer experiments revealed that when mature T cells were adoptively transferred into *Rag2^-/-^* mice (alone or in combination with B cells), this caused a pronounced tightening of the BBB (**Fig. 2A** (cortex) **Fig S1A** (hippocampus), **Fig S1B** (cerebellum)) and blood-CSF barrier (**Fig. 2B**). Interestingly, the transfer of B cells alone did not restore the integrity of the barriers. Furthermore, antibody-mediated depletion experiments in wild type mice supported the crucial contribution of T cells to the sealing capacity of the brain barriers. Here, depletion of T cells in wild type mice significantly increased the permeability of both the BBB (**Fig. 2C** (cortex), **Fig S1C** (hippocampus), **Fig S1D** (cerebellum)) and blood-CSF barrier (**Fig. 2D**). In contrast, depletion of B cells did not have a significant impact on barrier function. As expected, when both T and B cells were depleted, the increase in barrier permeability was comparable to that observed with T cell depletion alone.

These findings underscore the importance of T cells in preserving brain barrier integrity and suggest that their presence is essential for maintaining the protective function of the barriers. Because of the brain-protective role that has recently been assigned to Tregs^28^, we were prompted to test whether they are responsible for the maintenance of brain barrier integrity. Indeed, adoptive transfer of CD4^+^CD25^+^ T cells into *Rag2^-/-^* mice resulted in a significant decrease in permeability of the BBB (**Fig. 2E** (cortex), **Fig S1E** (hippocampus) and **Fig S1F** (cerebellum)) and blood-CSF barrier (**Fig. 2F**). Additionally, antibody-mediated depletion of Tregs in wild type mice revealed that the leakiness of both the BBB (**Fig. 2G** (cortex) **Fig S1G** (hippocampus), **Fig S1H** (cerebellum)) and the blood-CSF barrier (**Fig. 2H**) were significantly increased compared to wild type mice injected with an isotype control. These findings reveal a novel, non-canonical function of Tregs in maintaining brain barrier integrity.

**Figure 2.**
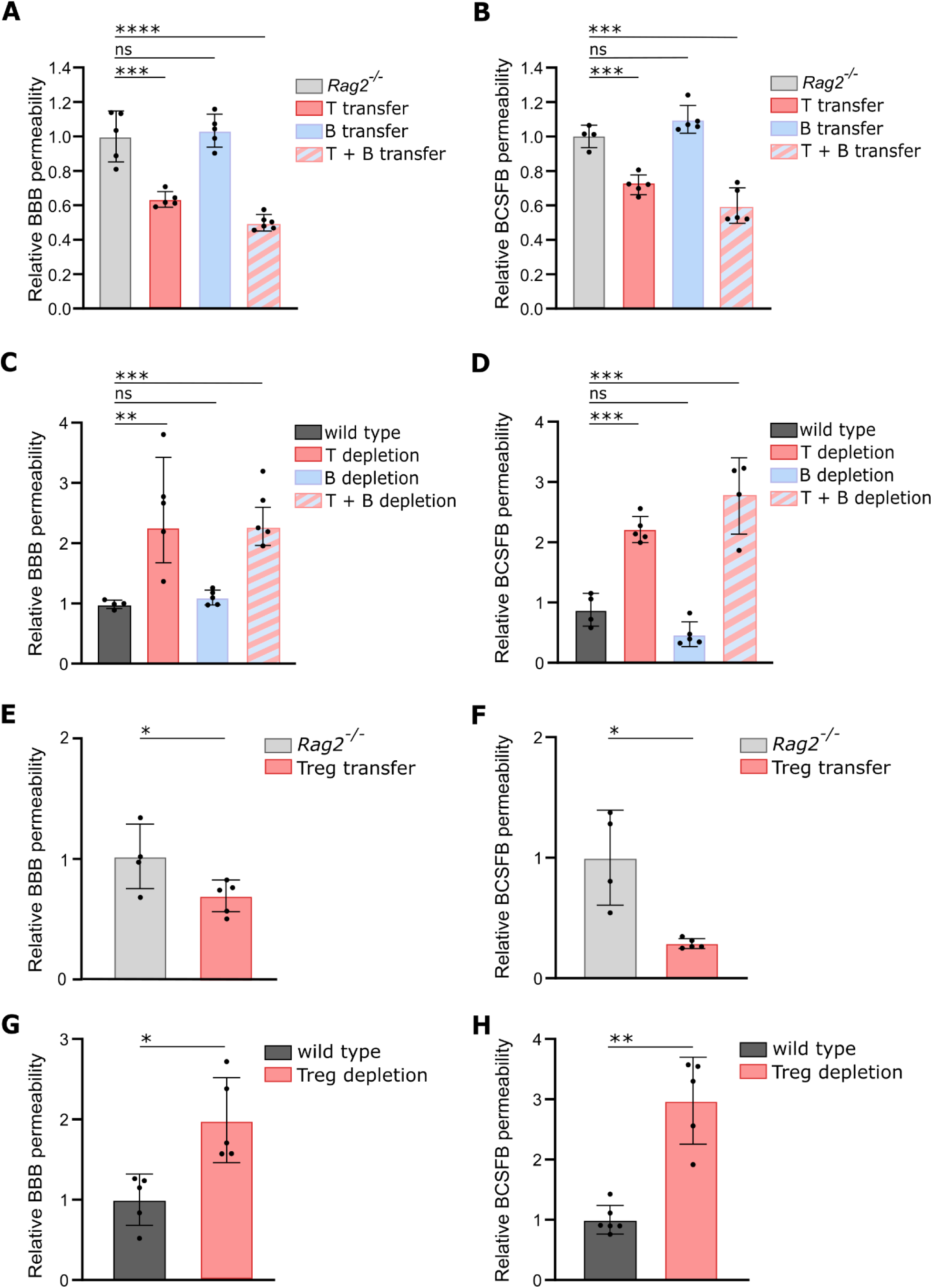
The integrity of the brain barriers relies on the presence or absence of Treg cells. Barrier permeability was determined by measuring the 4 kDa fluorescein isothiocyanate (FITC)-dextran leakage into the brain parenchyma and CSF 15 min after intravenous injection. (**A-B**) Relative BBB permeability in the cortex (A) and at the blood-CSF barrier (B) in wild type compared to *Rag2^-/-^*mice. T and/or B cells were isolated from the spleen of wild type donor mice and adoptively transferred to *Rag2^-/-^*acceptor mice (n=5). (**C-D**) In wild type mice, antibodies against T cells were intraperitoneally (i.p.) injected every 3 days for 15 days. Antibodies against B cells were i.p. injected every 5 days for 15 days. Wild type mice injected with non-reactive isotype antibodies were used as control. The relative permeability at the BBB in the cortex (C) and at the blood-CSF barrier (D) compared to wild type mice is shown. (n=5) **(E-F)** Regulatory T (Treg) cells were isolated from the spleen of wild type donor mice and adoptively transferred to *Rag2^-/-^* acceptor mice. The relative BBB permeability in cortex (E) and blood-CSF barrier permeability (F) compared to *Rag2^-/-^* mice is shown. (n=4-5) (**G-H**) In wild type mice, antibodies against Treg cells were i.p. injected every 3 days for 15 days. Wild type mice injected with non-reactive isotype antibodies were used as control. The relative BBB permeability in cortex (**F**) and blood-CSF barrier permeability (**G**) compared to wild type mice is shown. (n=5) The graphs are shown as the mean ± SEM and the datapoints are biological replicates. Statistical significance was determined by one-way ANOVA and Dunnett’s multiple comparisons test; * p < 0.05; ** p < 0.01; *** p < 0.01, **** p < 0.0001, ns: non-significant. BBB: blood-brain barrier; BCSFB: blood-cerebrospinal fluid barrier

**Figure S1.**
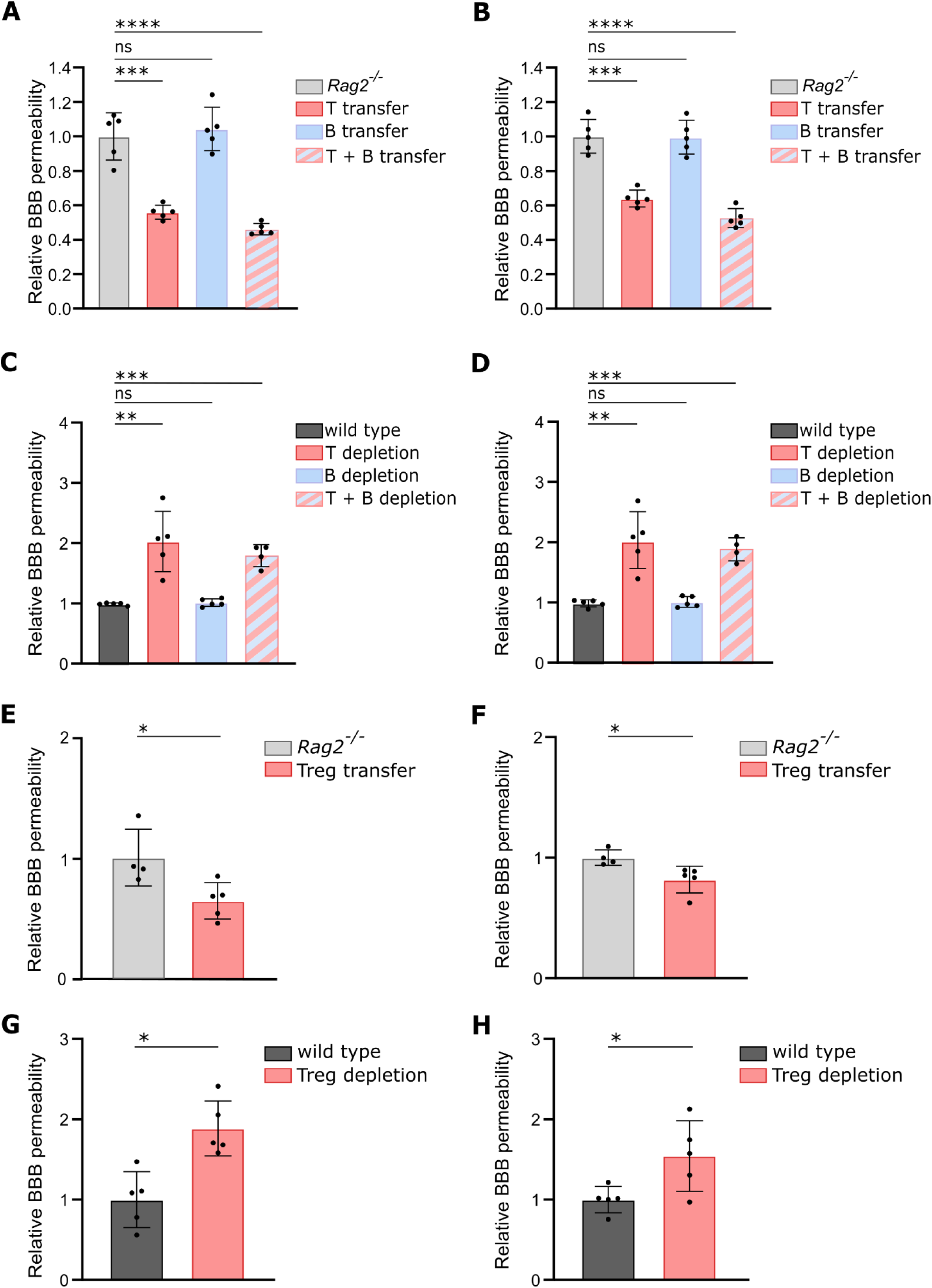
The integrity of the blood brain barriers relies on the presence or absence of T cells. Barrier permeability was determined by measuring the 4 kDa fluorescein isothiocyanate (FITC)-dextran leakage into the brain parenchyma 15 min after intravenous (i.v.) injection. (**A-B**) Relative BBB permeability in the hippocampus **(A)** and cerebellum (**B**) compared to *Rag2^-/-^* mice is shown. T and/or B cells were isolated from the spleen of wild type donor mice and adoptively transferred to *Rag2^-/-^* acceptor mice (n=5). (**C-D**) In wild type mice, antibodies against T cells were intraperitoneally (i.p.) injected every 3 days for 15 days. Antibodies against B cells were i.p. injected every 5 days for 15 days. Wild type mice injected with non-reactive isotype antibodies were used as control. The relative BBB permeability in the hippocampus (**C**) and cerebellum (**D**) compared to wild type mice is shown (n=5). **(E-F)** Regulatory T (Treg) cells were isolated from the spleen of wild type donor mice and adoptively transferred to *Rag2^-/-^*acceptor mice. The relative BBB permeability in the hippocampus (**E**) and the cortex (**F**) compared to *Rag2^-/-^*mice is shown (n=4-5). (**G-H**) In wild type mice, antibodies against Treg cells were i.p. injected every 3 days for 15 days. Wild type mice injected with non-reactive isotype antibodies were used as control. The relative BBB permeability in the hippocampus (**G**) and the cerebellum (**H**) compared to wild type mice is shown (n=5). The graphs are shown as the mean ± SEM and the datapoints are biological replicates. Statistical significance was determined by one-way ANOVA and Dunnett’s multiple comparisons test; * p < 0.05; ** p < 0.01; *** p < 0.001, **** p < 0.0001, ns: non-significant. BBB: blood-brain barrier; BCSFB: blood-cerebrospinal fluid barrier.

### IL-34, a cytokine produced by human Tregs, strengthens brain barriers by increasing ZO-1 expression

Having identified a novel brain barrier-protective effect of Tregs, we were prompted to reveal the underlying mechanism through which Tregs tighten the brain barriers. IL-34, which has been described as a Treg-specific cytokine implicated in their immunoregulatory functions^29^, is an interleukin that is functionally similar to macrophage colony-stimulating factor (M-CSF). Its functions are just starting to be uncovered, and up to now it has mainly been recognized for its role in stimulating survival, differentiation, migration and function of various myeloid mononuclear cells and macrophages^30^. Interestingly, previous research by Jin *et al*^31^ has hinted at the reparative role of IL-34 in BBB endothelial cells. Here, using flow cytometry, we could confirm IL-34 expression in Tregs (CD4^+^FoxP3^+^), although it was not exclusive to Tregs, as other CD4^+^ T cells (CD4^+^FoxP3^-^; non-Tregs) also showed IL-34 expression. Indeed, our findings revealed a significant increase in the percentage of IL-34 expressing Tregs (**Fig. 3A**) and non-Tregs (**Fig. 3B**), as well as an elevation in the median fluorescence intensity (MFI) of IL-34 in both Tregs (**Fig. 3C**) and non-Tregs (**Fig. 3D**) following stimulation. Nonetheless, Tregs exhibited a significantly higher percentage of IL-34 expressing cells (**Fig. 3E**) and MFI of IL-34 (**Fig. 3F**) compared to non-Tregs.

**Figure 3.**
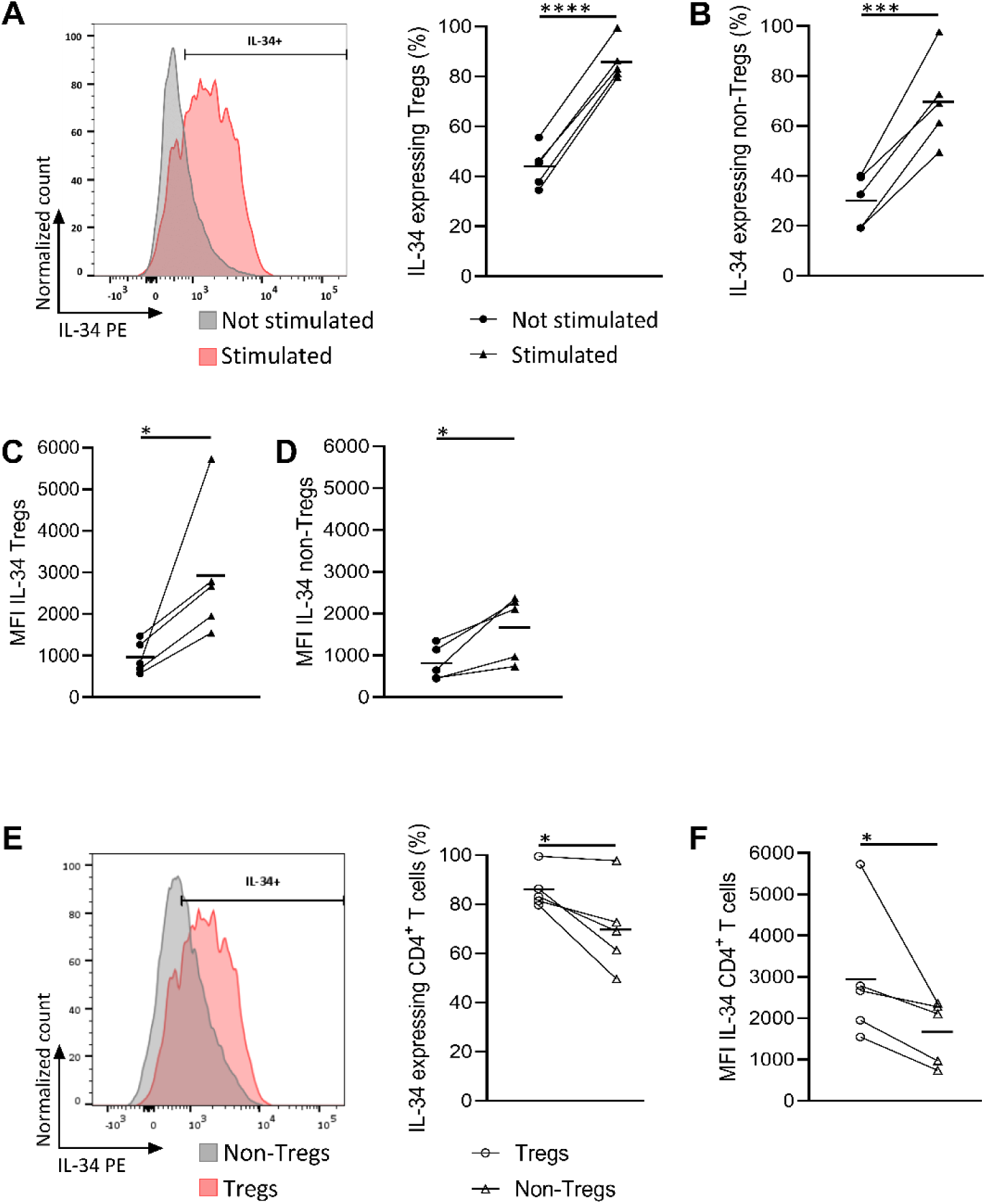
Human Tregs produce IL-34. CD4^+^ T cells were isolated from human blood and stimulated with or without Treg suppression inspector beads (72h) followed by flow cytometric analysis for IL-34 expression within regulatory T cells (Treg) (CD4^+^FoxP3^+^) or non-Tregs (CD4^+^FoxP3^-^). (**A-B**) Percentage of IL-34 expressing Tregs (A) and non-Tregs (B) with or without stimulation. Showing a representative plot of the normalized count of Tregs with and without stimulation (n=5). (Paired t-test) (**C-D**) Median fluorescence intensity (MFI) of IL-34 in Tregs (**C**) and non-Tregs (**D**) with or without stimulation (n=5). (Wilcoxon matched-pairs signed rank test) (**E-F**) Percentage of IL-34 expressing cells (**E**) (paired t-test) or MFI of IL-34 (**F**) (Wilcoxon matched-pairs signed rank test) within stimulated Tregs or non-Tregs (n=5). Histograms are normalized to the maximum count of the represented conditions. Values are paired per donor and horizontal bar represent group mean. Gating strategy is depicted in Fig. S3. Statistical significance was determined by paired t-test or Wilcoxon matched-pairs signed rank test. * p < 0.05; ** p < 0.01; *** p < 0.001; **** p < 0.0001

**Figure S2.**
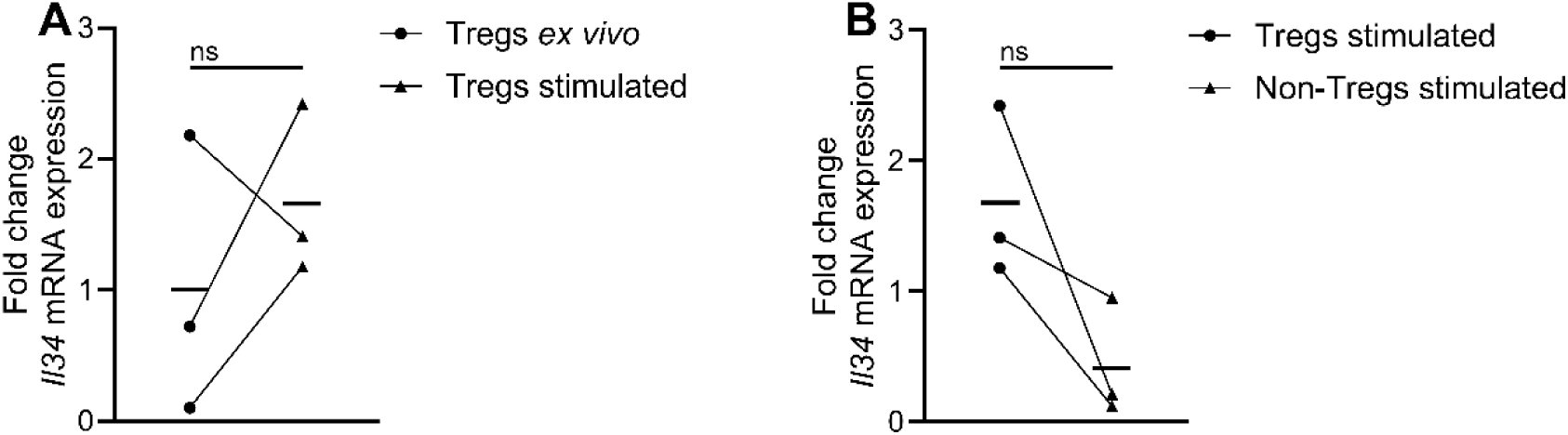
Human Tregs express *Il34* mRNA. Regulatory T cells (Tregs, CD4^+^CD25^hi^) and non-Tregs (CD4^+^CD25^-^) were isolated from human blood and collected immediately (*ex vivo*) or stimulated with Treg suppression inspector beads (72h). (**A**) Fold change of *Il34* mRNA expression in *ex vivo* and stimulated Tregs (n=3). (**B**) Fold change of *Il34* mRNA expression in stimulated Tregs and non-Tregs. (n=3). Values are paired per donor and horizontal bar represent group mean. Statistical significance was determined by Wilcoxon matched-pairs signed rank test. ns; non-significant.

**Figure S3.**
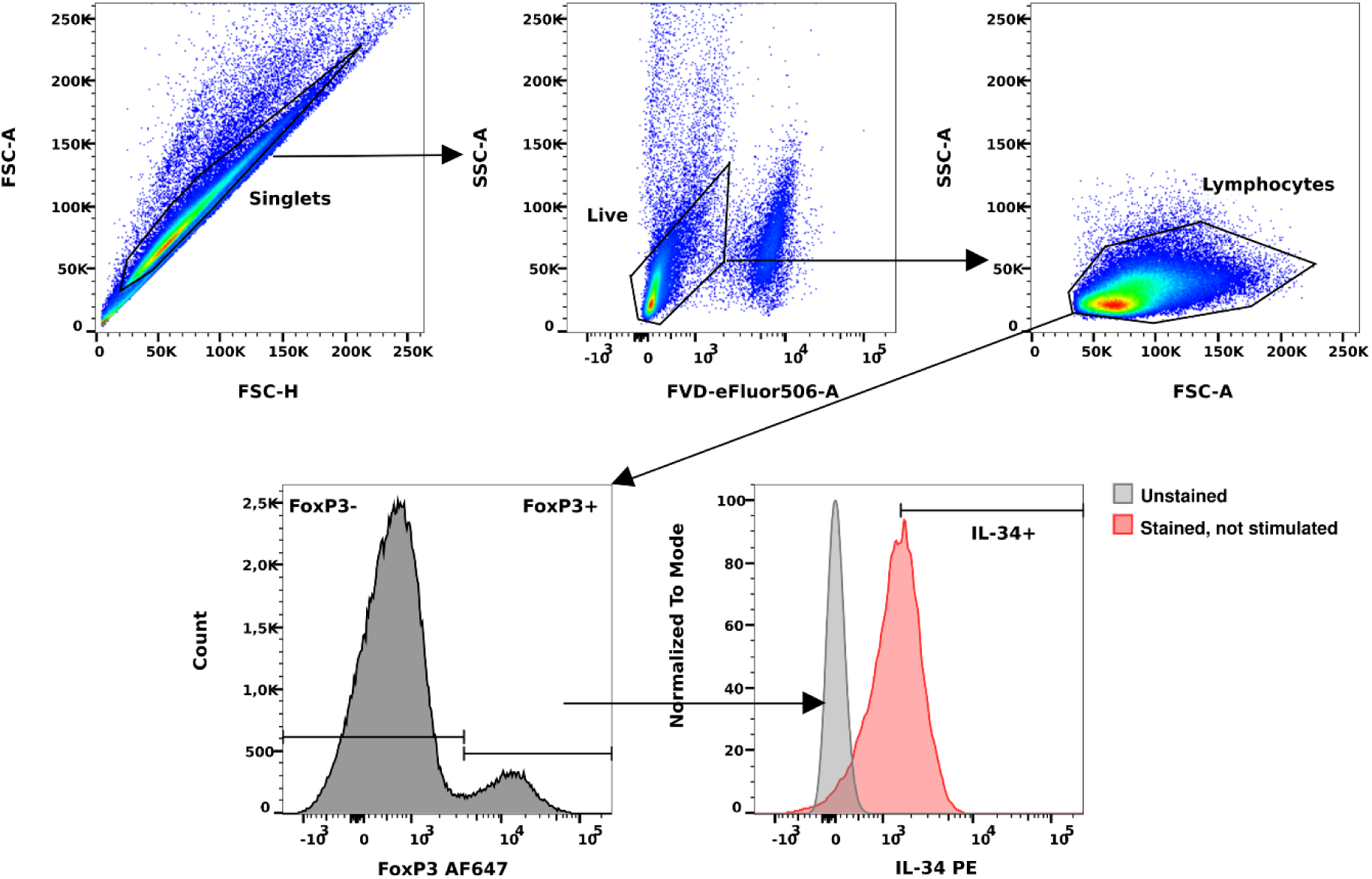
Gating strategy for demonstrating IL-34 expression in human Tregs (FoxP3^+^) and non-Tregs (FoxP3^-^). Human CD4^+^ T cells were isolated using magnetic activated cell sorting (MACS) and stimulated or not stimulated using Treg Suppression Inspector Beads followed by flow cytometric analysis. Gating strategy and representative plots are shown of non-stimulated CD4 T cells.

These results were validated at mRNA level, wherein stimulation led to a trend in increased *Il34* mRNA expression in Tregs (**Fig. S2**), with Tregs demonstrating higher expression compared to non-Tregs (**Fig. S2**). In summary, our findings demonstrate that Tregs express IL-34 and upregulate it upon stimulation, and that this IL-34 response is more pronounced compared to other CD4^+^ T cells.

Next, we studied whether IL-34 can directly influence brain barrier integrity. To accomplish this, we carried out *in vitro* diffusion assays to measure permeability of the brain barriers to 4 kDa FITC-dextran (FD). In a first set-up, primary BBB endothelial cells (BBB-ECs) and ChP epithelial (CPE) cells from *Rag2*^-/-^ mice were used (**Fig. 4A-B**). Interestingly, once a stable TEER was reached, the addition of IL-34 resulted in a tighter barrier, reflected by reduced 4 kDa FD leakage across the primary EC and CPE cell layers. In a next set-up, primary BBB-ECs and CPE cells from wild type mice were used (**Fig. 4D**). Here, after reaching a stable TEER, we stimulated the cells with the pro-inflammatory trigger LPS, resulting in increased leakage of 4 kDa FD across both brain barriers (**Fig. 4C**). When IL-34 was added simultaneously, the LPS-induced loss in barrier integrity was prevented (**Fig. 4D**). In addition, similar experiments were performed using immortalized cell lines, more specifically murine ChP epithelial (ImmCPE) and brain endothelial cells (Bend.3) (**Fig. 4E**). Similarly, after stimulation with LPS, an increased leakage of 4 kDa FD across both brain barriers was induced. When IL-34 was added simultaneously, the LPS-induced loss in barrier integrity was prevented. Next, we analysed whether IL-34 could influence ZO-1 expression. Remarkably, our results reveal that when IL-34 was added at the same time as the LPS stimulus, it prevented the LPS-induced reduction in ZO-1 expression both in Bend.3 (**Fig. 4F**) and immCPE cells (**Fig. 4G**).

**Figure 4.**
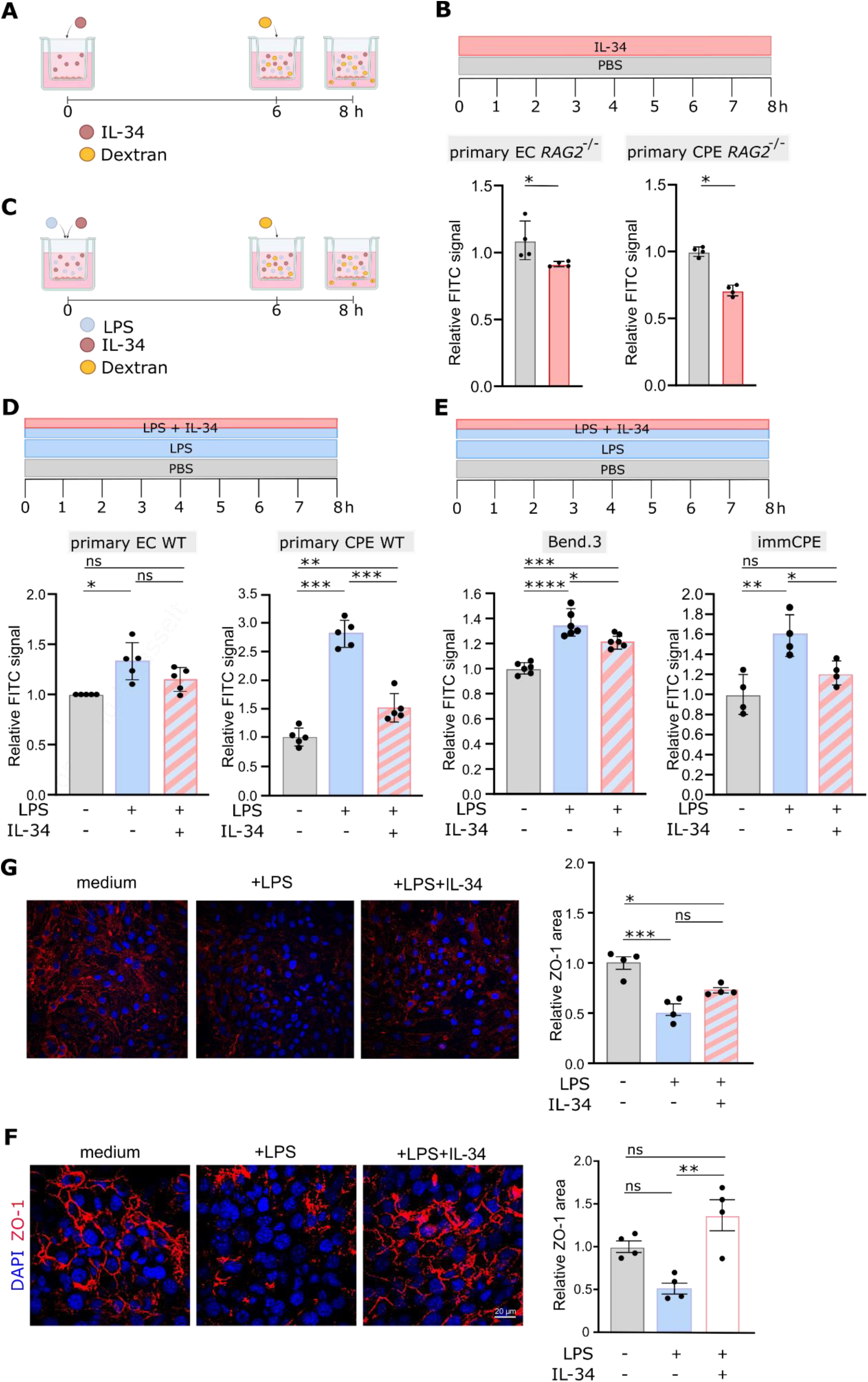
*In vitro* transwell studies show that IL-34 positively effects brain barrier tightness and ZO-1 expression level. (**A**) Schematic representation of transwell assay used in (B). Upon reaching a constant trans-epithelial electrical resistance (TEER), cells were stimulated with medium or medium containing IL-34 for 8 h. At the 6h timepoint, 4 kDa fluorescein isothiocyanate (FITC)-dextran was added. (**B**) Primary choroid plexus epithelial (CPE) cells and brain endothelial cells (ECs) were isolated from *Rag2^-/-^* mice and seeded in transwell system. Relative leakage of 4 kDa fluorescein isothiocyanate (FITC)-dextran compared to PBS treated cells is shown (n=4). (**C**) Schematic representation of transwell assay used in (D-G). Upon reaching a constant TEER, cells were stimulated with LPS or LPS and IL-34 for 8 h. After 6h 4 kDa FITC-dextran was added. (**D**) Primary CPE cells and brain ECs were isolated from wild type mice. Relative leakage compared to PBS treated cells is shown (n=5). (**E**) Murine immortalized CPE cells (ImmCPE) and murine immortalized EC cells (Bend.3) were seeded. Relative leakage compared to PBS treated cells is shown (n=4-6). (**F-G**) Representative images of ZO-1 expression in confluent Bend.3 cell layer (F) and confluent ImmCPE cell layer (G). Graphs represent quantification of the ZO-1 signal. The graphs are shown as the mean ± SEM and the datapoints are biological replicates. Images are representative for 4 biological replicates. Statistical significance was determined by Mann-Whitney test to compare to groups or by one-way ANOVA and Dunnett’s multiple comparisons test to compare multiple groups; * p < 0,05; **p < 0.01; *** p < 0.001; **** p < 0.001; ns: non-significant.

Altogether, these findings highlight the ability of IL-34 to tighten brain barriers, at least in part through increasing ZO-1 expression.

### IL-34 expression by human Tregs is altered during MS and AD

Various neurological disorders, including MS^32^ and AD^33^, exhibit early disruption of the BBB and blood-CSF barrier. To understand whether IL-34 expression in Tregs is also altered in these diseases, we analyzed blood samples of MS and AD patients, as well as subjects with MCI, a precursor stage of the Alzheimer’s disease continuum, preceding overt dementia. Interestingly, these analyses revealed that untreated RR-MS patients show a decreased frequency of IL-34 expressing Tregs (**Fig. 5A**) in the blood, compared to matched healthy donors (HD), and this was not the case for IL-34 expressing non-Tregs (**Fig. 5B**). Additionally, the MFI of IL-34 was reduced in Tregs derived from RR-MS patients compared to HD (**Fig. 5C**). Next to patients with RR-MS, AD and MCI patients also exhibited a lower frequency of IL-34 expressing Tregs (**Fig. 5D, G**), but not in non-Tregs (**Fig. 5E, H**), in the blood compared to matched HD. Here, the MFI of IL-34 within Tregs was not altered due to disease (**Fig. 5F,I**). These findings underscore the potential translational significance of Treg-derived IL-34 as a brain barrier-repairing molecule in various neurological diseases.

**Figure 5.**
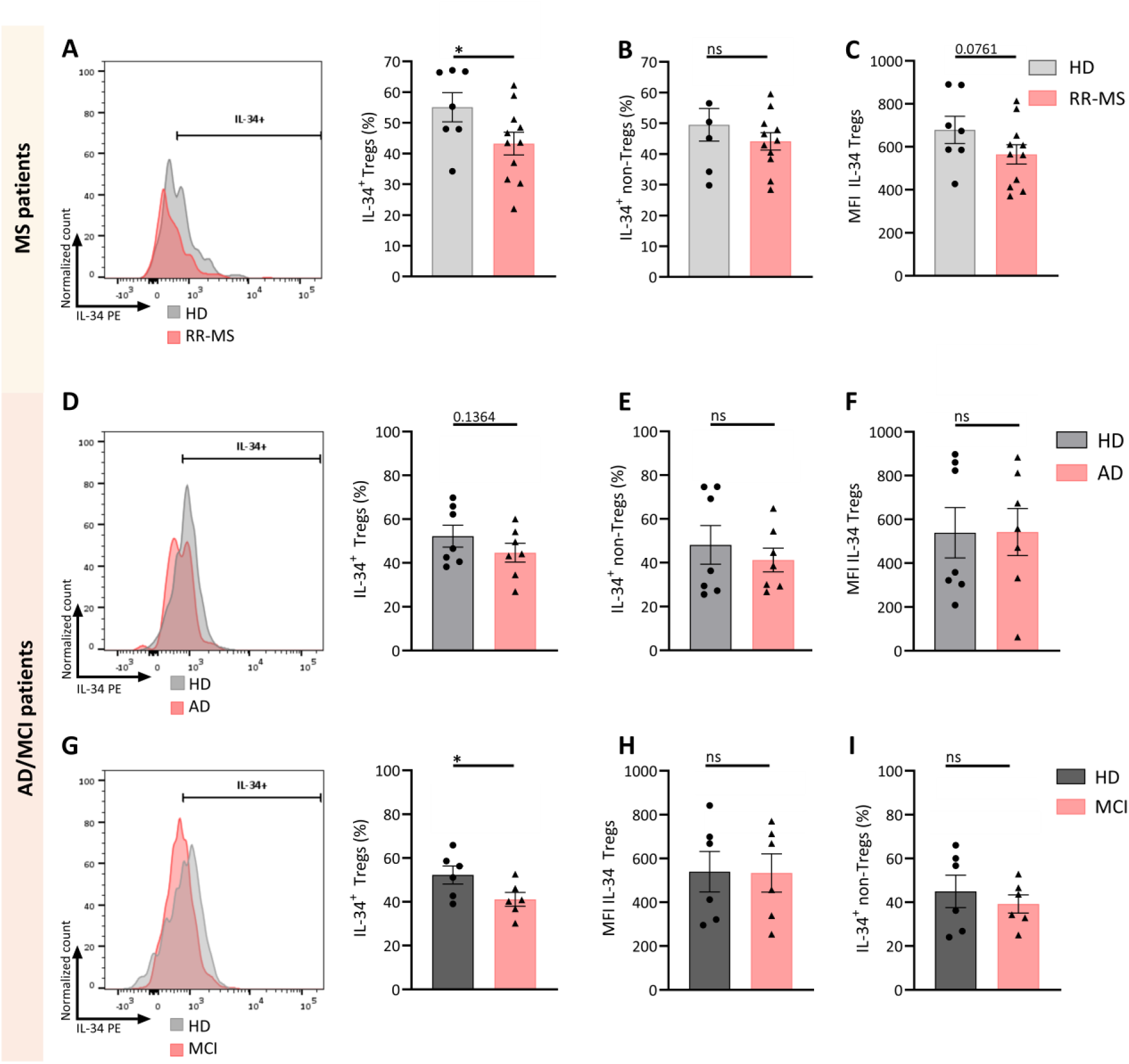
IL-34 expression is reduced in Tregs from MS, AD and MCI patients in contrast to non-Tregs. **(A-G)** Frozen PBMCs or whole blood from healthy donors (HD) (n = 7), untreated relapsing-remitting multiple sclerosis (RR-MS,n=11), Alzheimer’s disease (AD,n=7) or mild cognitive impairment (MCI,n=6) patients were thawed and stimulated for 4h with PMA, CaI and Protein Transport Inhibitor Cocktail. Cells were stained and analysed for flow cytometry. Representative plot of the normalized regulatory T cell (Treg) count and percentages of IL-34^+^ within Tregs (CD4^+^FoxP3^+^) of RR-MS (**A**), AD (**D**) and MCI (**G**) patients, and within non-Tregs (CD4^+^FoxP3^-^) of RR-MS (**B**), AD (**E**) and MCI (**H**) patients. Median fluorescence intensity (MFI) of IL-34 in Tregs of RR-MS (**C**), AD (**F**) and MCI (**I**) patients. Data are represented as mean ± SEM and the datapoints are biological replicates. Gating strategy is depicted in Fig. S4. Statistical significance was determined by unpaired t-test (MS graphs) or Mann-Whitney test (AD/MCI graphs); *p<0.05, ns: non-significant.

**Figure S4.**
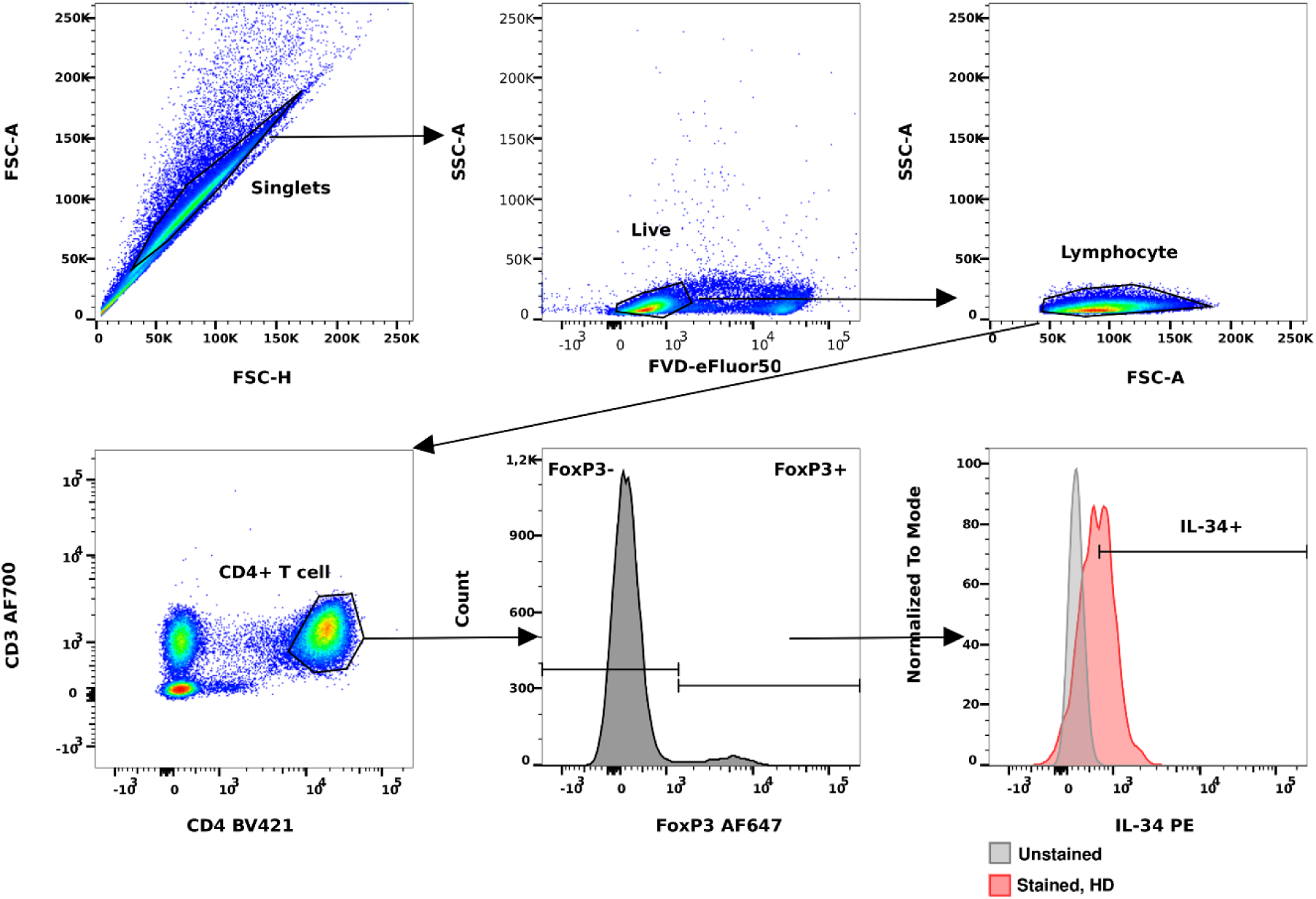
Gating strategy of IL-34 in patient and healthy donor samples. Frozen PBMCs and whole blood were analysed using flow cytometry. Gating strategy and representative plots are shown of healthy donor (HD) PBMCs.

### IL-34 administration improves brain barrier integrity in a murine model for MS and AD

Following these novel findings, we aimed to explore the therapeutic potential of IL-34 in bolstering brain barriers in murine models of neurodegenerative disorders. Notably, in experimental autoimmune encephalomyelitis (EAE) mice, a widely used model for MS, we observed a decrease in IL-34 levels in the blood at 16 and 21 days post-induction (**Fig. 6A**). Subsequently, we investigated whether augmenting the IL-34 concentration in the bloodstream through thrice-daily systemic administration for three days could ameliorate barrier permeability in EAE mice (**Fig. 6B**). As depicted in **Fig. 6C**, elevating IL-34 levels in the blood at peak of disease, led to a reduction in both BBB and blood-CSF barrier leakage.

**Figure 6.**
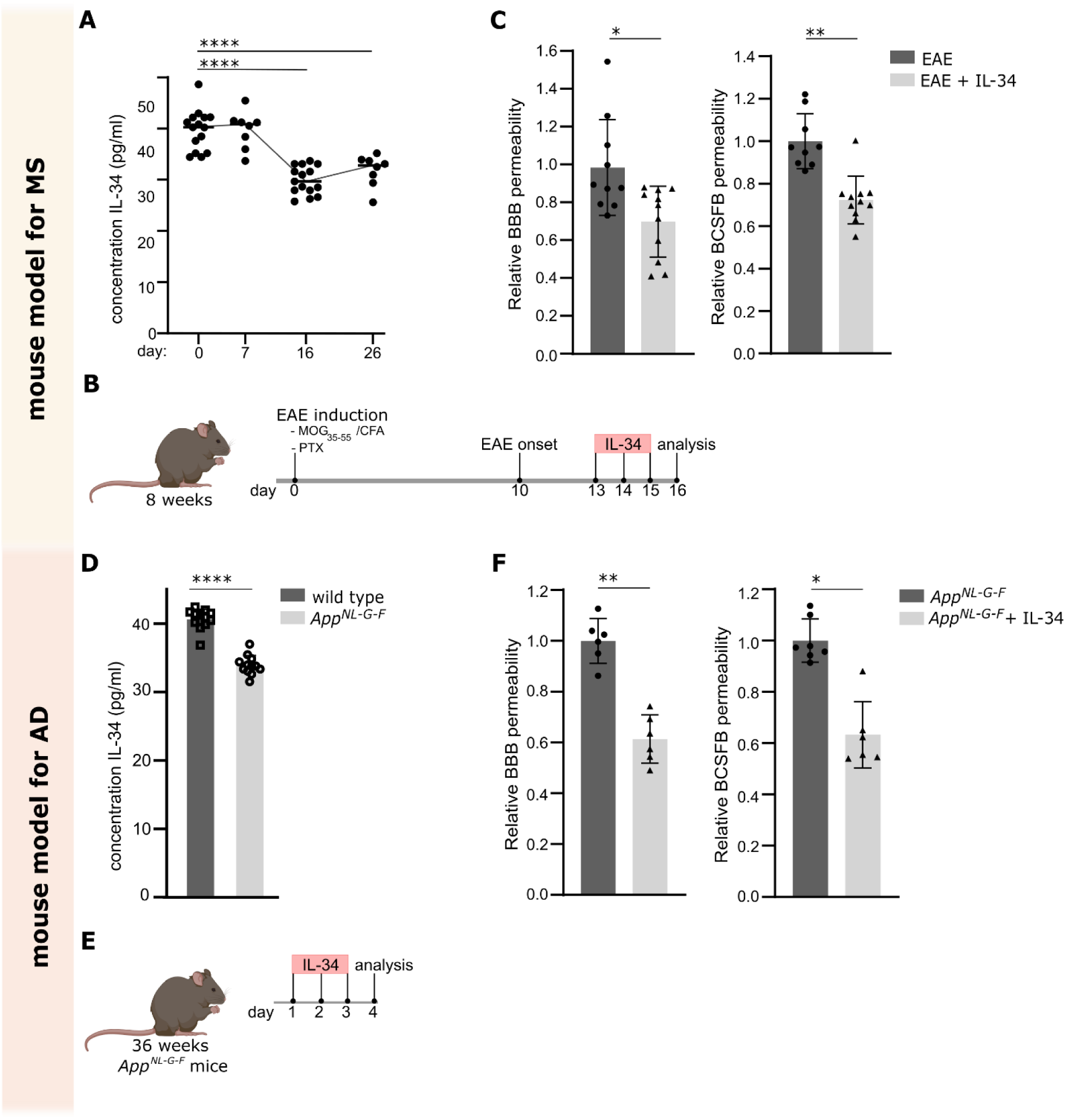
IL-34 administration tightens brain barriers in murine models of MS and AD. (**A**) IL-34 levels in the blood of mice during the course of experimental autoimmune encephalomyelitis (EAE) (n=8 or 15). (**B**) Schematic representation of treatment scheme. IL-34 was intravenously (i.v.) administered 3 times a day for 3 days starting on day 13 post-EAE induction. On day 16, barrier leakage was determined by measuring the 4 kDa fluorescein isothiocyanate (FITC)-dextran leakage into the brain parenchyma 15 min after i.v. injection. (**C**) Relative BBB permeability in the hippocampus and relative blood-CSF barrier permeability is shown (n=9 or 11). (**D**) IL-34 levels in the blood of age matched wild type and *App^NL-G-F^* mice at 36 weeks (n=14). (**E**) Schematic representation of treatment scheme. IL-34 was i.v. administered 3 times a day for 3 days. On day 4, barrier leakage was determined by measuring the 4 kDa FITC-dextran leakage into the brain parenchyma 15 min after i.v. injection. (**F**) Relative BBB permeability in the hippocampus and relative blood-CSF barrier permeability is shown (n=6). The graphs are shown as the mean ± SEM and the datapoints are biological replicates. Statistical significance was determined by student t-test or by one-way ANOVA and Dunnett’s multiple comparisons test to compare multiple groups; * p < 0.05; **p < 0.01; **** p < 0.0001.

By using a murine model for AD, namely *App^NL-G-F^* mice, we additionally showed lower levels of IL-34 detected in the blood of these diseased mice (**Fig. 6D**). By increasing the blood concentration via thrice-daily systemic administration for three days of IL-34 (**Fig. 6E**), a reduction in leakage across both brain barriers was observed (**Fig. 6F**).

Combined, these experiments highlight the therapeutic potential of an IL-34 based therapy for improving brain barrier integrity in neurodegenerative disorders.

Finally, we set out to unravel the molecular mechanism observed upon IL-34 treatment *in vivo*. Thereto, we administered IL-34 i.v. to endotoxemic mice every 2h as depicted in **Fig. 7A**, followed by quantification of BBB and blood-CSF barrier leakage. Remarkably, our results demonstrate that repetitive systemic administration of IL-34 was able to protect against inflammation-induced BBB leakage (**Fig. 7B** and **Fig. S5**) and blood-CSF barrier leakage (**Fig. 7D**). These findings align with our earlier *in vitro* data, reinforcing the notion that IL-34 plays a crucial role in regulating brain barrier integrity. In addition, and in agreement with our *in vitro* results, this was associated with restoration of ZO-1 levels that showed severe disturbance upon endotoxemia (**Fig. 7C** and **7E**). These results underpin the therapeutic value of IL-34 treatment to protect brain barriers in neuro-inflammatory conditions.

**Figure 7.**
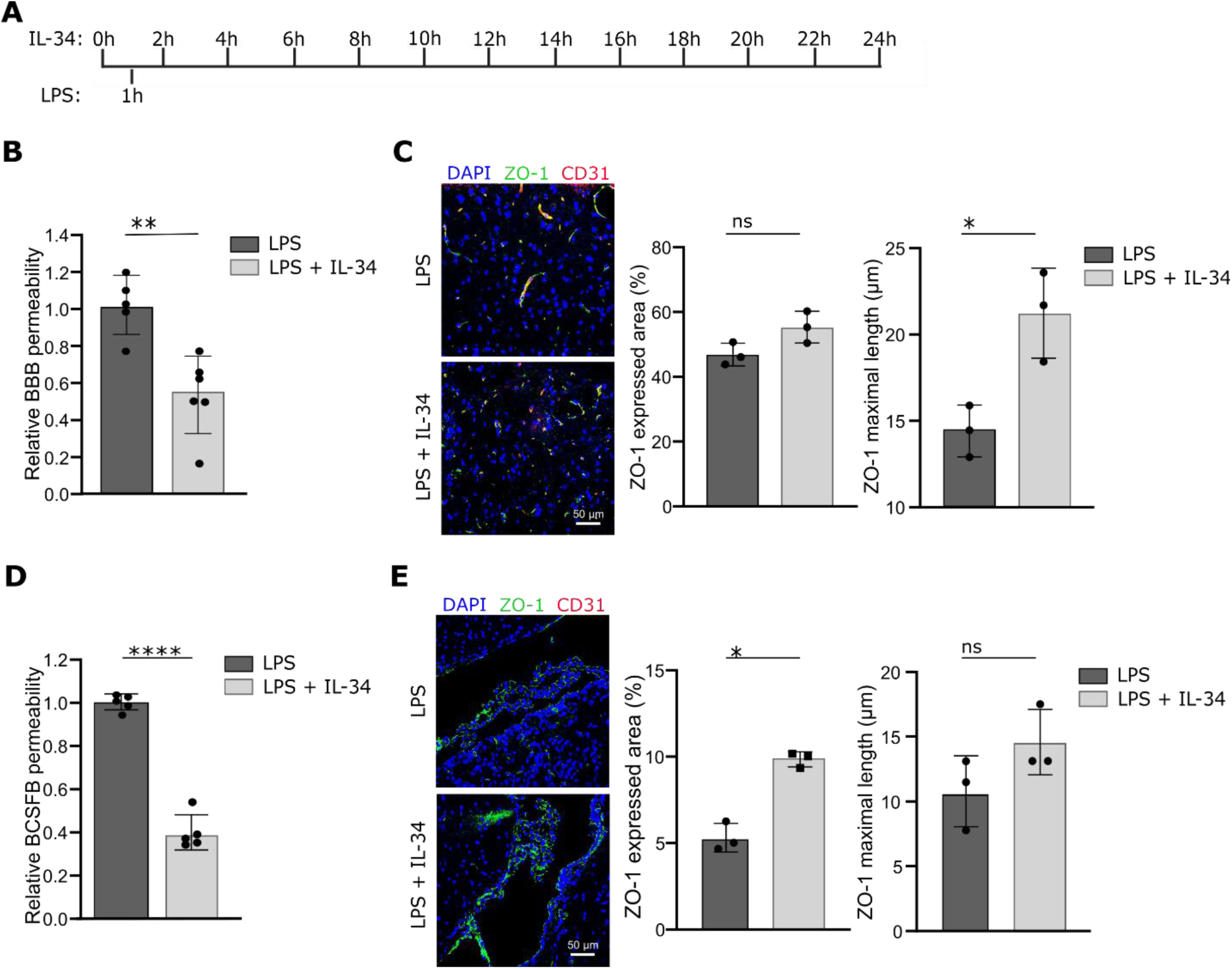
IL-34 prevents LPS-induced loss of barrier integrity *in vivo* and this is associated with maintenance of ZO-TJ levels. (**A**) Experimental set-up: Endotoxemia was induced via intraperitoneal injection of LPS (2.5 mg/kg). Every h IL-34 (75 pg) was administered intravenously (i.v.), followed by analysis 24 h later. (**B** and **D**) Barrier permeability was determined by measuring the level of 4 kDa fluorescein isothiocyanate (FITC)-dextran 15 min after i.v. injection. The relative BBB permeability in the cortex (**B**) and CSF (**D**) compared to wild type mice is shown. (**C** and **E**) Representative images of ZO-1 expression at the BBB (**C**) and blood-CSF barrier (**E**). Graphs represent quantification of the ZO-1 signal. Scale bar: 50 µm. Images are representative for 4 biological replicates. The graphs are shown as the mean ± SEM and the datapoints are biological replicates. Statistical significance was determined by Mann-Whitney test; * p < 0,05; ** p < 0,01; **** p < 0.0001.

**Figure S5.**
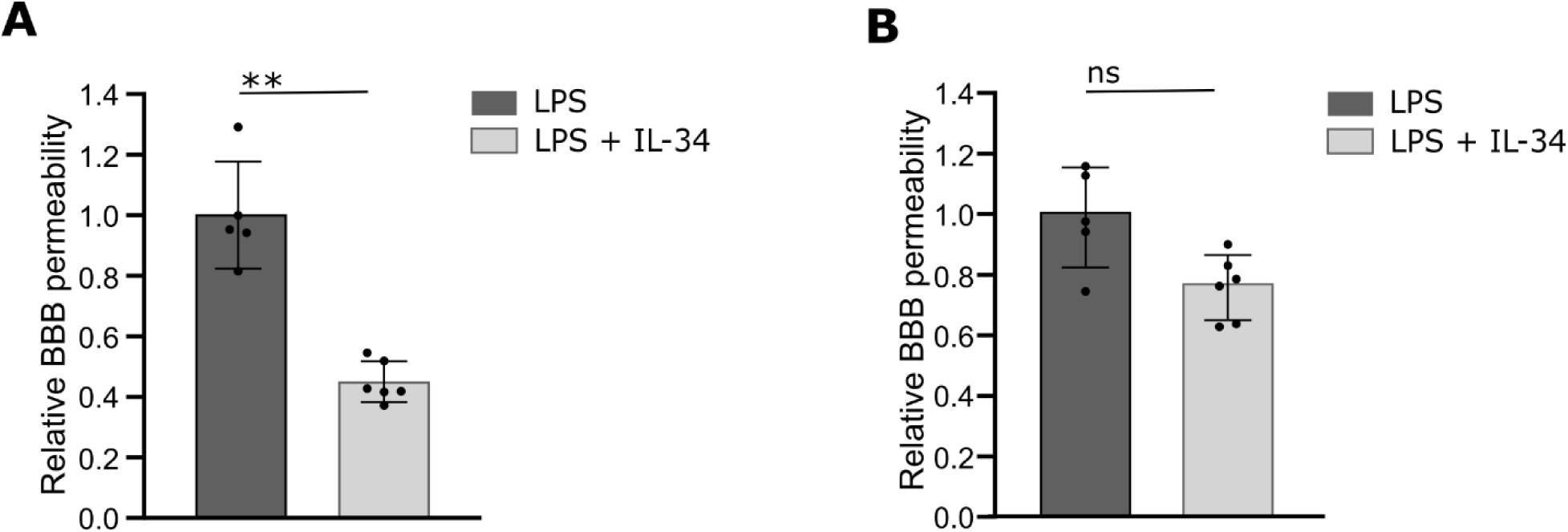
IL-34 prevents LPS-induced loss of blood-brain barrier integrity in hippocampus and cerebellum. Endotoxemia was induced via intraperitoneal injection of LPS (2.5 mg/kg). During 24 h after the LPS injection, every 2 h IL-34 (75 pg) was administered intravenously (i.v.), followed by analysis 24 h later. Barrier permeability was determined by measuring the level of 4 kDa fluorescein isothiocyanate (FITC)-dextran 15 min after i.v. injection. The relative BBB permeability in the hippocampus (**A**) and cerebellum (**B**) compared to endotoxemia mice that received IL-34 is shown. The graphs are shown as the mean ± SEM and the datapoints are biological replicates. Statistical significance was determined by Mann-Whitney test; ** p < 0,01; ns: non-significant.

## Discussion

In addition to their well-established immunosuppressive role, Tregs have recently been discovered to possess non-canonical regenerative functions, such as promoting neural stem cell proliferation and suppressing neurotoxic astrogliosis^14,15,34^. Furthermore, an *in vivo* study in 2 rodent models of ischemic stroke indicated that Treg therapy indirectly protects the blood-brain barrier (BBB) through the suppression of the production of matrix metalloprotease-9 derived from peripheral neutrophil ^35,36^. Notably, a common observation in neurological disorders like multiple sclerosis (MS) and Alzheimer’s disease (AD) is a reduction in Treg functionality and/or numbers^37^. The compromised integrity of brain barriers is a crucial factor in the pathogenesis of MS and is increasingly recognized as a significant contributor to the progression of AD^38,39^. However, the direct link between Treg dysfunction or reduction and compromised brain barrier integrity in these diseases has remained unclear so far. Here, we are the first to identify a novel non-canonical function of Tregs, demonstrating their pivotal role in maintaining the sealing capacity of both the BBB and the blood-cerebrospinal fluid (CSF) barrier. Through a series of adoptive transfer and depletion experiments, we have now unveiled the direct involvement of Tregs in preserving brain barrier integrity.

In addition, we elucidated the mechanism underlying this newfound non-canonical function of Tregs. Our findings reveal that human Tregs produce the cytokine IL-34, which alone has the capability to enhance the sealing capacity of both the BBB and the blood-CSF barrier *in vitro* and *in vivo*. IL-34, primarily expressed in the brain by neurons and in the skin by keratinocytes, plays a crucial role in immune tolerance post-organ transplantation^29,40,41^. Within the brain, IL-34 has been shown to sustain microglia and suggested BBB integrity by upregulating tight junction (TJ) expression in endothelial cells (ECs) following inflammation^40,41,31^. Expanding upon these findings, our study employed primary brain ECs, choroid plexus epithelial (CPE) cells, and immortalized murine cell lines in *in vitro* diffusion assays, revealing that direct application of IL-34 is sufficient to enhance barrier tightness. Additionally, we further elucidate the mechanism of action by showing that improvements upon IL-34 treatment involve increased expression of TJ protein ZO-1.

To validate these direct effects of IL-34 *in vivo*, we administered IL-34 to endotoxemic mice at physiological levels, effectively enhancing brain barrier integrity by increased TJ expression. However, our *in vivo* set-up does not exclude the possibility that the observed effects on brain barrier sealing capacity are solely attributable to the direct influence of IL-34. In the CNS, IL-34 produced by neurons binds to colony stimulating factor 1 receptor (CSF-1R) on microglia, promoting their differentiation and proliferation^40^. Although microglia play a role in the neurovascular unit of the BBB, studies indicate that microglia depletion has minimal impact on BBB integrity^42,43^. Besides microglia, leptomeningeal and perivascular macrophages also express CSF-1R, suggesting that IL-34 treatment may affect their function, thereby contributing to the observed *in vivo* effects on brain barrier integrity. Additionally, macrophage populations at the blood-CSF barrier, including choroid plexus (ChP) macrophages at the apical epithelial surface and stromal macrophages in the stromal space, may also contribute to improved barrier integrity upon IL-34 stimulation. However, we pin this *in vivo* effect down to the same mechanism as seen *in vitro*, the increased expression levels of TJ ZO-1.

Strikingly, we were able to demonstrate the therapeutic potential of an IL-34-based therapy for neuro-inflammatory disorders characterized by impaired brain barriers. Our investigation reveals a decrease in the frequency of IL-34 expressing Tregs in patients with relapsing-remitting MS (RR-MS) compared to healthy controls. Previous studies examining IL-34 levels in MS have yielded conflicting results. Serum analysis showed no significant differences between RR-MS and control groups^44^, consistent with our observations of IL-34 expression in the overall immune cell population. However, conflicting results were reported in the CSF. One study observed downregulation in MS patients compared to healthy controls^45^, while another found no differences^46^. In contrast, a study with a control group of pseudotumor cerebri patients showed increased IL-34 levels in MS. However, this control group is not suitable due to elevated CSF volume, potentially diluting cytokine concentrations^47^. These diverse findings underscore the complex role of IL-34 in physiological and disease conditions, with no published reports investigating its specific expression in Tregs during disease states. In a previous study, mice lacking IL-34 exhibited accelerated development of experimental autoimmune encephalomyelitis (EAE) without a significant increase in disease severity^48^. Our findings suggest that decreased IL-34 cytokine levels in the blood of EAE mice are associated with disease progression. This reduction in IL-34, and consequently, the loss of its protective effect, may lead to a more rapid breakdown of the barriers, consistent with our data on IL-34’s protective effect on brain barrier integrity. While the systemic administration of IL-34 at physiological levels significantly enhanced brain barrier integrity, it did not translate into improved disease progression in r EAE mice. This lack of efficacy might be due to the brief duration of IL-34 treatment, lasting only 3 days, with a subsequent follow-up period of just 1 day, which may have been insufficient to detect any therapeutic effects.

The pathology of AD involves compromised barrier integrity, enabling the infiltration of inflammatory cells. This initiates a vicious cycle, where the accumulation of inflammatory cells and Aβ further worsens brain barrier function, thereby exacerbating AD pathology^49^. Consequently, sealing the brain barriers presents a promising therapeutic approach for AD. A recent publication identified a genetic variant in *Il34* as risk factor for AD development^50^, while another study has previously speculated on the contribution of IL-34 to AD pathogenesis, particularly in relation to processes associated with the brain barriers^50^. Here, we have uncovered that, similar to MS patients, AD patients have a lower frequency of IL-34 expressing Tregs in their blood. Even in MCI as the early stage of dementia, we see similar changes, suggesting early involvement of Tregs in AD pathology. Supporting this notion, we demonstrated decreased IL-34 cytokine levels in the blood compared to age-matched control mice. Notably, by systemically administering IL-34 at physiological levels in *App*^NL-G-F^ mice, we observed a significant improvement in brain barrier integrity. Throughout our study, we observed that IL-34 has a more pronounced impact on the blood-CSF barrier compared to the BBB. Both barriers are comprised of polarized barrier cells with specific TJs and adherens junctions (AJs), resulting in distinct characteristics in their apical and basal membranes. This distinction may extend to differences in the expression profiles of membrane receptors. Our *in vitro* and *in vivo* experiments indicate that IL-34 primarily influences the cells from the blood side of the brain barriers, rather than from the CSF or brain parenchymal side. Therefore, in our transwell studies, IL-34 was administered to the blood side, while *in vivo* experiments utilized intravenous injections of IL-34. Importantly, accumulating evidence suggests that during inflammation, T cells tend to accumulate predominantly in the perivascular spaces rather than infiltrating the brain parenchyma^51^.

Our study highlights the critical role of Tregs in maintaining the integrity of both the BBB and the blood-CSF barrier, revealing a novel non-canonical function of Tregs and identifying IL-34 as a crucial cytokine in this process. Mechanistically, IL-34 modulates the strength of the brain barriers by influencing the TJ protein ZO-1. Importantly, our data demonstrate that IL-34 treatment effectively restores the integrity of brain barriers in mouse models of AD and MS. These ground-breaking findings provide valuable insights into the intricate relationship between Tregs, IL-34, and brain barrier integrity, presenting new possibilities for therapeutic strategies aimed at maintaining CNS homeostasis and alleviating dysfunction of brain barriers in neurological disorders.

## Material and methods

### Mice

The different mouse colonies were maintained in specific pathogen free (SPF) conditions. Mice were kept in individually ventilated cages under a 10-h dark/14-h light cycle with free access to food and water. Both male and female mice were used and randomly allocated to experimental groups, generating gender balanced groups. All animal studies were conducted in compliance with governmental and EU guidelines for the care and use of laboratory animals and were approved by the ethical committee of the Faculty of Sciences, Ghent University, Belgium (EC2022-118 and EC2023-085).

### Mouse models

*Rag2^-/-^* mice are deficient in the *RAG2* gene and fail to generate mature T and B lymphocytes (RRID:IMSR_JAX:008449, Jackson Laboratory). *App^NL-G-F^* mice carry the Arctic, Swedish, and Iberian mutations as described in Saito *et al*^52^ and are used as murine model for Alzheimer’s disease (AD). Experimental autoimmune encephalomyelitis (EAE) as murine model for multiple sclerosis (MS) mice were generated by immunization of 8 to 15-week-old C57BL/6 wild type female mice with the MOG_35-55_/CFA Emulsion PTX Hooke Kit (EK-2110; Hooke Laboratories) following the manufacturer’s guidelines. Briefly, mice were subcutaneously injected with an emulsion of MOG_35-55_ in complete Freund’s adjuvant (CFA) two times (total of 200 µg/mouse), followed by administration of pertussis toxin (PTX, 121 ng/mouse) in Dulbecco’s phosphate buffered saline (D-PBS, 14190-094; Gibco), first on the day of immunization and then again 24 h later. For all experiments, age and sex matched C57BL/6 J mice were used as a control.

### Primary mouse cells

Primary mouse choroid plexus epithelial (CPE) cells were isolated from P2-P7 pups as previously described ^53^. Choroid plexus tissue was isolated from the lateral and fourth ventricles, pooled and digested with pronase (537088, Sigma-Aldrich) for 7 min. For monolayer cultures, cells were plated on Transwell polyester inserts (pore size, 0.4 μm; surface area, 33.6 mm^2^; 3470, Corning) coated with laminin (L2020-1mg, Sigma-Aldrich). Cells were grown in Dulbecco’s Modified Eagle Medium (DMEM, 41965-039; Gibco) DMEM medium supplemented with 10% fetal bovine serum (FBS), 2 mM L-Glutamine (Gibco), 1% penicillin/streptomycin at 37°C and 5% CO_2_ for 1 to 2 weeks until their trans-epithelial electrical resistance (TEER) values reached a plateau.

Primary mouse brain microvascular endothelial cells, were isolated from four to six-week-old mice with a C57BL/6 background as described before^54^. Brains were dissected followed by removal of the meninges, mincing and homogenization. The tissue was enzymatically digested, and cells were collected by using a 33% continuous Percoll gradient (17089101, cytiva). The cells were cultured in DMEM (D5796, Sigma) containing 20% FBS (Biowest), 1 ng/ml FGF (F0291), 100 μg/ml heparin (H3149), 1.4 μM hydrocortisone (H0888, all Merck) and 0.5% penicillin/streptomycin (P4333, Sigma) at 37°C and 5 % CO^2^, and cells were plated on 10 μg/ml collagen type IV (234154, Merck)-coated Thincerts (3 µm, translucent, 24-well plate, 662631, Greiner Bio-One). To purify the cell culture, medium was supplemented with 10 μg/ml puromycin (A11138-03, Sigma-Aldrich) for 48 h, 4 μg/ml puromycin for 24 h and no puromycin for the remaining culturing time. TEER values were measured to evaluate growth.

### Cell lines

The murine immortalized brain endothelial *cell* line (*bEnd*.*3*) (Watanabe *et al*, 2013) and the murine immortalized choroid plexus epithelium cell line immCPE (Pauwels *et al*, 2022) were used to study the blood-brain barrier (BBB) and blood-cerebrospinal fluid (CSF) barrier *in vitro*. The bEnd.3 cells were cultured in DMEM supplemented with L-glutamine (BE17-605 F; Lonza), Na-pyruvate (S-8636; Sigma), penicillin-streptomycin (P4333; 100x solution Sigma) and 10% FBS. These cells were cultured in DMEM/F12 medium (11320033; Gibco) supplemented with respectively 10% or 20% FCS, in laminin (L2020; Sigma-Aldrich)-coated flasks. Media were further supplemented with non-essential amino acids (M-7145; 100x solution Sigma), L-glutamine (BE17-605 F; Lonza), Na-pyruvate (S-8636; Sigma-Aldrich), penicillin-streptomycin (P4333; 100x solution Sigma-Aldrich). All cells were cultured at 37 °C and 5% CO2.

### Quantification of BBB and blood-CSF barrier permeability

The brain barrier permeability was evaluated according to the methods described before^55^. Shortly, 4 kDa FITC-labelled dextran (FD4-1G; Sigma-Aldrich) was injected i.v. 15 min before CSF collection. CSF was obtained from the fourth ventricle via cisterna magna puncture using needles made from borosilicate glass capillary tubes (B100-75-15; Sutter Instruments) (Balusu *et al*, 2016; Vandenbroucke *et al*, 2012). Next, mice were anaesthetized with a lethal dose of ketamine/xylazine (100 mg/kg ketamine; 20 mg/kg xylazine) and perfused with D-PBS/heparin (0.2% heparin, H-3125; Sigma-Aldrich), afterwards, brain regions were isolated and snap frozen in liquid nitrogen. CSF was diluted 100-fold in sterile D-PBS and the blood-CSF barrier leakage was measured by quantifying fluorescence at λ_ex_/λ_em_ = 485/520 nm on the FLUOstar Omega (BMG Labtech). Brain samples were finely cut and incubated overnight at 37°C in formamide (47671; Sigma-Aldrich) while shaking in the dark. The next day, the supernatant was collected after centrifugation at 400 g-forces (g) for 7 min. The degree of BBB leakage was determined by measuring fluorescence at λ_ex_/λ_em_ = 485/520 nm on the FLUOstar Omega.

### Visualization of BBB and blood-CSF barrier permeability

4kDa lysine fixable dextran (D7135; Thermo Fisher) was i.v. injected. 15 min later mice were anesthetized with ketamine/xylazine (100 mg/kg ketamine; 20 mg/kg xylazine in D-PBS) and decapitated. The brain was dissected from the skull and fixed overnight in 4% PFA at 4°C. To prepare for cryo-preservation, samples were sequentially incubated in 15% and 30% sucrose (27,483,294; VWR) in D-PBS solutions at 4 °C. Next, tissue samples were embedded in NEG-50 (6502; Prosan bvba) and stored at -80 °C. These samples were cut into 20 μm cryosections (CryoStar NX70, Thermo Scientific). For immunofluorescence staining, sections were permeabilized in 0.3% PBST (PBS containing 0.3% Triton X-100). Following being blocked with goat immunomix (GIM) (5% goat serum in 0.3% PBST) at 21 °C for 1h, sections were incubated with CD31 (553370; BD Pharmingen) in GIM at 4°C overnight. After washing with PBS, sections were stained with fluorophore-conjugated secondary antibodies in 0.1% PBST at 21 °C for 1.5h. Next, samples were counterstained with DAPI reagent (Sigma-Aldrich, 1:1000 in PBS) and the sections were mounted.

### Immunohistochemistry

For immunostainings on brain sections, mice were transcardially perfused with ice-cold 4% PFA in PBS. Subsequently, brains were carefully separated from the skull and split into two halves (mid-sagittal). The right hemispheres were embedded in Frozen Section Medium (Thermo Scientific) immediately in cryomolds (Sakura) that were frozen on dry ice and stored at –80 °C until further use. The brains were cut into 20 μm cryosections (CryoStar NX70, Thermo Scientific). These brain samples were cut from the sagittal superior sinus towards the border of the cerebral hemisphere. Sections were collected serially for TJ staining when the choroid plexus was initially present. One section per mouse was analysed. For immunofluorescence staining, sections were permeabilized in PBS containing 0.3-0.5% Triton X-100. Following being blocked with goat immunomix (GIM) (5% goat serum, 0.1% BSA, 0.3-0.5% Triton X-100 in PBS) at 21 °C for 1h, sections were incubated with primary antibodies in GIM at 4°C overnight. After washing with PBS, sections were stained with fluorophore-conjugated secondary antibodies in PBS or PBS containing 0.1% Triton X-100 at 21 °C for 1-1.5h. Counterstaining was done with Hoechst reagent (Sigma-Aldrich, 1:1000 in PBS). Primary antibody ZO-1 (61-7300; Thermo Scientific) and CD31 (DIA-310; Dianova) were used. For immunostainings on cells, cells were fixed with 2% PFA for 20 min on ice. Next, cells were washed three times with PBS and permeabilized with 0.1% PBST for 10 min on ice. Samples were washed with blocking buffer (1% BSA in PBS) and incubated with primary antibodies (diluted in blocking buffer) for 2h at 21 °C or overnight at 4°C. After washing with PBS, cells were stained with fluorophore-conjugated secondary antibodies in PBS for 1 at 21 °C. Next, samples were counterstained with Hoechst and the sections were mounted. A Zeiss LSM780 confocal microscope or Zeiss Axioscan Z.1 was used for imaging.

### Adoptive transfer experiments

The adoptive transfer of T and B cells was conducted on a 1:1 donor-to-acceptor mouse basis according to sex. Spleens were isolated from wild type mice and transferred to PBS with 0.5% BSA. Next, spleens were passed through a cell strainer (70 μm;734-0003; BD Falcon) to make a single-cell suspension. The cells were precipitated by centrifugation for 6 min at 300 g and 4°C. After discarding the supernatant, red blood cells were removed using Ammonium-Chloride-Potassium (ACK) lysing buffer (10-548E; Lonza Bioscience). The cell pellet was resuspended in separation buffer (PBS, pH 7.2, 0.5% BSA, 2 mM ethylenediaminetetraacetic acid (EDTA)). Dead cells were stained with Trypan Blue (0.4%, 15250061; Gibco) and live cells were counted using a hematocytometer. T, B and Treg cells were further isolated by negative selection using the mouse Pan T cell isolation kit II (130-095-130; Miltenyi Biotec), the mouse Pan B cell isolation kit II (130-104-443; Miltenyi Biotec) and the mouse CD4^+^CD25^+^ Regulatory T Cell Isolation Kit (130-091-041; Miltenyi Biotec) following the manufacturer’s guidelines. After 7 days, barrier integrity was determined as described above.

### Antibody-mediated depletion experiments

In wild type mice, T cells were depleted by intraperitoneal (i.p.) injections of 300 µg/mouse of InVivoPlus anti-mouse CD3ε antibody (BP0001-1; BioXcell) dissolved in PBS every 3 days for 15 days. B cells were depleted by i.p. injections of 300 µg/mouse of InVivoMAb anti-mouse B220 (BE0067; BioXcell) dissolved in PBS every 5 days for 15 days. Treg cells were depleted by i.p. injections of 300 µg/mouse of InVivoMAb anti-mouse CD25 (BE0012; BioXcell) dissolved in PBS every 3 days for 15 days. Control animals received i.p. injections of 300 µg/mouse InVivoMAb rat IgG2a isotype control (BE0089; BioXcell), InVivoMAb polyclonal rat IgG (BE0094; BioXcell) and InVivoMAb rat IgG1 isotype control, anti-horseradish peroxidase (BE0088; BioXcell) dissolved in PBS for respectively T, B and Treg cells. On day 16, barrier integrity was determined as described above.

### *In vitro* barrier function assessments

To identify the *in vitro* effect of IL-34 on brain barrier integrity, primary EC, primary CPE, bEnd.3 or ImmCPE cell lines were grown on transwell collagen-coated membrane 0.4um pore size inserts (CLS3495-24EA; Sigma-Aldrich) in a 24-well plate. Before transferring the cells to the inserts, dead cells were stained with Trypan Blue (0.4%, 15250061; Gibco) and live cells were counted using a hematocytometer. To each insert, 1,5 × 10^8^ cells in 300 µl were added, and 500 µl of medium was added to the basal compartment. Confluency was determined by measuring the transepithelial/transendothelial electrical resistance (TEER) using the EVOM2 epithelial volt/ohm meter (World Precision Instruments). When the cells reached confluency, the following conditions were applied: PBS for 6h (control), LPS (1 µg/mL, tlrl-pelps; InvivoGen) for 6h, LPS (1 µg/mL) together with IL-34 (50 ng/mL, 5195-ML-010; R&D systems) for 6h, and LPS (1 µg/mL) for 6h with the addition of IL-34 (50 ng/mL) after 4h. At the 6h timepoint, 4 kDa FITC-dextran (FD4-1G; Sigma-Aldrich) was added to the basal compartments of the 24-well plate. After 2h, samples were taken from the apical compartment, followed by measuring the fluorescence at λ_ex_/λ_em_ = 485/520 nm on the FLUOstar Omega (BMG Labtech).

### *In vivo* administration of IL-34

Recombinant mouse IL-34 (75 pg, 5195-ML-010; R&D systems) was i.v. injected every 2h for 24h. Wild type mice additionally received an LPS injection (2.5 mg/kg) at the 1h timepoint. At the 24h timepoint, the mice were sedated, and the barrier integrity was determined as described above. EAE and *App^NL-G-F^* mice were i.v. injected 9 times spread over 3 days with recombinant mouse IL-34 (74,16 pg, 5195-ML-010; R&D systems) or with sterile D-PBS as control.

### Plasma isolation

Blood was taken from EAE and *App^NL-G-F^* mice by heart punction and centrifuged at 1300 g and 4°C for 10 min. Supernatant was taken and centrifuged again at 2400 g and 4°C for 15 min.

### Measurement of IL-34 by ELISA

To determine the level of IL-34 in the plasma *App^NL-G-F^*mice, 35-week-old mice and age-matched wild type controls were used. For EAE, mice of 7 days, 16 days, and 26 days after immunization with MOG_35-55_ were used. Naïve EAE mice were used as control. The level of IL-34 in the plasma of EAE and *App^NL-G-F^* mice was determined using the LEGEND MAX^TM^ Mouse IL-34 ELISA kit (439107; BioLegend) following the manufacturer’s guidelines.

### Human samples

PBMC freezings from MS patients and healthy donors (HD) and whole blood freezings from AD and MCI patients were derived from the University Biobank Limburg (UBiLim). Patient information is summarized in **Table 1** (RR-MS) and **Table 2** (AD and MCI). This study was approved by the Medical Ethical Committee of Hasselt university and Medical Ethical Committee of Ziekenhuis Oost-Limburg (ZOL). Informed consent was obtained from all study subjects or when applicable from their caregivers.

**Table 1:** Donor characteristics of RR-MS patients and healthy donors.

|  | HD | RR-MS |
| --- | --- | --- |
| Sex (M/F) | 2/5 | 2/9 |
| Age ± SEM | 30.8 ± 1.818 | 32.8 ± 1.995 |
| EDSS ± SEM | NA | 1.364 ± 0.234 |
| Treatment | NA | Untreated |

EDSS: expanded disability status scale; F: female; M: male; NA: not applicable; RR-MS: relapsing-remitting MS; SEM: standard error of the mean.

**Table 2:** Donor characteristics of AD and MCI patients and healthy donors.

|  | HD (AD) | AD | HD (MCI) | MCI |
| --- | --- | --- | --- | --- |
| Sex (M/F) | 6/1 | 6/1 | 3/3 | 3/3 |
| Age ± SEM | 70.71 ± 2.18 | 70.57 ± 1.94 | 64 ± 3.71 | 64.33 ± 3.71 |

HD: healthy donor; AD: Alzheimer's disease; MCI: mild cognitive impairment; F: female; M: male; SEM: standard error of the mean.

### Flow cytometry of IL-34 expression

PBMCs were isolated from whole blood by the ficoll density gradient method (CL5020) and CD4^+^ T cells were selected using the human CD4^+^ T cell isolation kit (130-096-533, Miltenyi Biotec) according to manufacturer’s instructions. All CD4^+^ T cells were stimulated or not using human Treg Suppression Inspector beads (130-092-909; Miltenyi Biotec) in X-VIVO 15 (02-060F; Lonza) with 5% FBS (Gibco). After 72h stimulation, cells were stained with the Fixable Viability Dye eFluor 506 (65-0866-14; eBioscience) followed by a staining with FoxP3 AF647 (320113; Biolegend) and IL-34 PE (IC5265P; R&D systems) antibodies. Cells were fixed and permeabilized using the FOXP3/Transcription factor staining buffer kit (00-5523-00; Thermo Fisher Scientific) following manufacturer’s instructions.

PBMCs and whole blood derived from RR-MS, AD, MCI and HD samples were cryopreserved in liquid nitrogen. After thawing, the cells were stimulated with 200 ng/ml phorbol 12-myristate 13-acetate (PMA; P1585; Merck), 100 ng/ml calcium ionomycin (CaI; 56092-82-1; Merck) and Protein Transport Inhibitor Cocktail (00-4980-93; Invitrogen) for 4h at 37 °C. Next, cells were stained using the Fixable Viability Dye eFluor 506 (65-0866-14; eBioscience) and following antibodies: CD25 BB515 (567318; BD Biosciences)/Kiravia Blue (356144; Biolegend), FoxP3 AF647 (320113), CD3 AF700 (300324), CD4 BV421 (300531; all Biolegend) and IL-34 PE (IC5265P; R&D systems). The FOXP3/Transcription factor staining buffer kit (00-5523-00; Thermo Fisher Scientific) was used for fixation. All samples were acquired on LSRFortessa (BD Biosciences) and analysed using FlowJo (version 10.8.1) (BD Biosciences). Gating strategies are depicted in **Fig. S3** and **S4**.

### Quantitative PCR

To investigate the gene expression level of *Il34* in (non-)stimulated Tregs and non-Tregs, PBMCs were isolated using the ficoll density gradient method followed using human CD25 MicroBeads II (130-092-983, Miltenyi Biotec) for positive MACS selection. CD25^+^ cells were further used for sorting using a FACSAria Fusion (BD Biosciences) using the following antibodies: CD4 AF700 (300526), CD25 Kiravia blue 520 (356144) and CD127 BV421 (351310; all Biolegend). The sorting strategy was used as shown before^56^. CD25^-^ cells underwent further selection for CD4 using the human CD4^+^ T cell isolation kit (130-096-533, Miltenyi Biotec) to establish the non-Treg population (CD4^+^CD25^-^). Cell pellets of Treg and non-Treg cells were collected and stored at -80°C directly after isolation or after 72h stimulation using human Treg Suppression Inspector beads (130-092-909, Miltenyi Biotec) in X-VIVO 15 (02-060F, Lonza) with 5% FBS (Gibco). Lysis of the cells was performed using RLT lysis buffer (Qiagen) with 1% β-mercapto-ethanol followed by RNA extraction using the RNeasy Plus Micro Kit (74034, Qiagen) according to manufacturer’s instructions. mRNA concentration was determined using the Nanodrop 2000/2000c Spectrophotometer (Thermo Fisher Scientific) and conversion of RNA to cDNA was done using qScript cDNA supermix (95048, Quanta Biosciences). Quantitative PCR (qPCR) was done using a SYBR Green PCR Master Mix (4309155, Applied Biosciences) followed by the reaction in Quantstudio 3 (Applied Biosystems). Primer sequences are listed in Table 3. Quantification of the gene expression levels was performed using the comparative Ct method and by normalizing to stable reference genes.

**Table 3:**
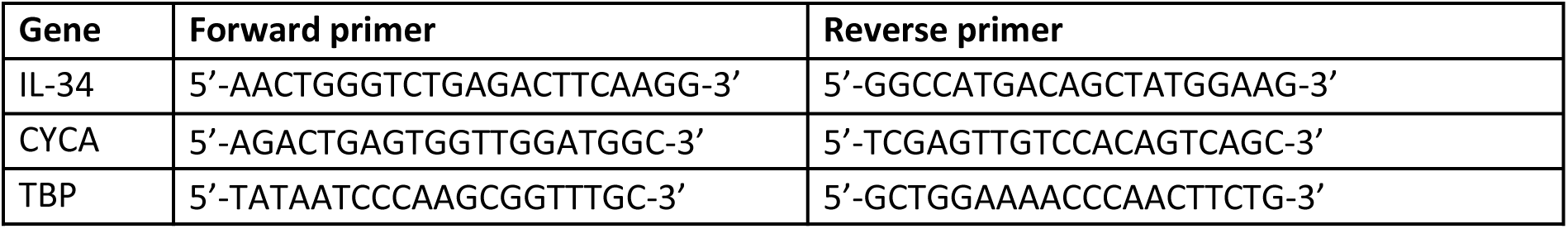
Primer sequences used for qPCR.

### Statistics

Data are shown as mean ± SEM. Two-tailed Student’s t-test for parametric data or Mann-Whitney for non-parametric data was used to compare two groups. One-way ANOVA with Dunnett’s multiple comparisons test was used to compare 3 or more groups. To compare 2 groups in paired samples, a paired t-test for parametric data and a Wilcoxon matched-paired signed rank test for non-parametric data was performed. GraphPad Prism 9.0 was used for statistical analysis. Differences were considered significant at p < 0.05.

## Acknowledgements

We thank the VIB Bioimaging Core Ghent and VIB flow cytometry Core Ghent for training, support and access to the instrument park. This work was supported by Research Foundation-Flanders (FWO) (1268823N,), Hasselt University (BOF23DOC22), Ghent Univeristy (BOF23/DOC/170), China Scholarship Council (CSC) (201808360194), Stichting voor Alzheimer onderzoek (SAO) (20200032), Charcot Foundation, Stichting MS Research (19-1064 MS), VLAIO (HBC.2022.0198).

